# CLEP: A Hybrid Data- and Knowledge-Driven Framework for Generating Patient Representations

**DOI:** 10.1101/2020.08.20.259226

**Authors:** Vinay Srinivas Bharadhwaj, Mehdi Ali, Colin Birkenbihl, Sarah Mubeen, Jens Lehmann, Martin Hofmann-Apitius, Charles Tapley Hoyt, Daniel Domingo-Fernández

## Abstract

As machine learning and artificial intelligence become more useful in the interpretation of biomedical data, their utility depends on the data used to train them. Due to the complexity and high dimensionality of biomedical data, there is a need for approaches that combine prior knowledge around known biological interactions with patient data. Here, we present CLEP, a novel approach that generates new patient representations by leveraging both prior knowledge and patient-level data. First, given a patient-level dataset and a knowledge graph containing relations across features that can be mapped to the dataset, CLEP incorporates patients into the knowledge graph as new nodes connected to their most characteristic features. Next, CLEP employs knowledge graph embedding models to generate new patient representations that can ultimately be used for a variety of downstream tasks, ranging from clustering to classification. We demonstrate how using new patient representations generated by CLEP significantly improves performance in classifying between patients and healthy controls for a variety of machine learning models, as compared to the use of the original transcriptomics data. Furthermore, we also show how incorporating patients into a knowledge graph can foster the interpretation and identification of biological features characteristic of a specific disease or patient subgroup. Finally, we released CLEP as an open source Python package together with examples and documentation.

## 1. Introduction

Recent advancements in machine learning (ML) and artificial intelligence (AI) methodologies have initiated a paradigm shift in bioinformatics. As new technologies have steadily generated large volumes of -*omics* data, AI methods have garnered great insights into human health and biology. With the availability of large-scale biological datasets, ML/AI are becoming highly relevant for biomedical applications, such as predictive modeling, patient stratification, and simulation [1]. However, despite the successful application of ML/AI in the biomedical domain, the datasets underlying the generation of models can play a far more crucial role in a given application than the complexity of the model itself. For example, in some cases, if data is predictive enough, simpler methods can outperform state-of-the-art ML/AI methods in prediction tasks [2, 3].

The development of novel high-throughput experimental techniques has led to a broad availability of biological data from multiple entity types [4]. In practice, integrating multiple data types can be advantageous, particularly in the context of complex diseases, where no single type of data can effectively explain the cause of dysfunction. However, biological datasets tend to be both inherently complex and noisy [5], making their integration challenging. Furthermore, biological data typically contain a far greater number of features than samples due to several factors, including a lack of available resources and obstacles in sample collection [6]. In failing to address these challenges and generating comprehensive representations of biological data, novel techniques can suffer in a range of analytic tasks.

One approach adopted by the systems biology community is in representing biological data in the form of networks. By doing so, multiple scales of biology can be represented, as can the relationships within and across these scales. These networks are also advantaged by their ability to integrate heterogeneous biological data types [7]. Generally, one can classify biological networks into two categories depending on the source of information used to generate them [8].

The first of these two classes of networks are constructed from biological data by using a variety of methodologies. For instance, co-expression networks can be generated from transcriptomics data to represent pairwise correlations between genes [9]. Another example are bayesian networks which can be used to model conditional interdependencies across heterogeneous biological entities [10]. While the majority of these methods transform biological data into networks comprising relations between the biological entities under study, other methods directly translate patient-level data into networks as an indicator of similarity between patients. These patient similarity networks closely link patients that are more similar to each other while less similar patients contain fewer close connections. Patient similarity networks have been successfully used to represent multimodal patient level-data for various classification and clustering tasks [11, 12, 13, 14, 15].

A second class of networks can be generated from prior knowledge of known interactions between biological entities. When sets of these interactions are assembled, they can be used to represent discrete biological networks. These networks can be referred to as knowledge graphs (KGs) when they comprise entities from various biological modalities and the complex interactions between them. However, despite their advantages, networks cannot be directly represented in vector space; thus, impeding their direct use by ML/AI techniques. This impediment has led to the development of methodologies designed to encode entities and relations in a KG into a latent feature space while preserving its structure. While these new representations can be used for entity disambiguation, clustering, and several downstream ML/AI tasks, they have been primarily used in the biomedical domain for link prediction tasks, including the prediction of side effects [16], disease-gene associations [17], and novel therapeutic targets [18]. However, until now, there have yet to be integrative approaches which incorporate both classes of these networks and leverage both from patient-level data and prior knowledge.

In this work, we introduce CLEP (CLinical Embedding of Patients), a hybrid data- and knowledge-driven framework that exploits patient-level data and incorporates this information into a KG. Once the KG has been generated by CLEP, the framework then drives the generation of novel patient representations through various knowledge graph embedding models (KGEMs). In building upon previous data-driven approaches [11, 15] which have demonstrated the advantages of representing patients as nodes in a network, we show that additionally integrating prior knowledge can make for more robusts analyses. We showcase our approach by generating new patient representations derived from two different transcriptomics datasets and a KG comprising heterogenous protein-protein interactions from multiple biological databases. Using the new patient representations, we find that performance on a panel of ML models trained to classify between patients and controls is significantly improved with respect to the use of original data. Furthermore, the flexibility of our approach makes it applicable to heterogenous multimodal biological datasets, enabling researchers to generate new patient representations by combining patient-level data with the prior knowledge contained in a KG. Finally, we have made CLEP available to the bioinformatics community as an open source Python package (https://github.com/hybrid-kg/clep) under the Apache 2.0 License.

## 2. Results

### 2.1. Framework Description

**Figure 1** illustrates each of the steps of the presented framework. The methodology requires a patient-level dataset and a KG. Patients are incorporated into the KG as new nodes and connected to features that most closely characterize a given patient. Once patients are embedded into the KG, KGEMs are then used to generate new patient representations which can subsequently be employed for a variety of downstream applications.

**Figure 1.**
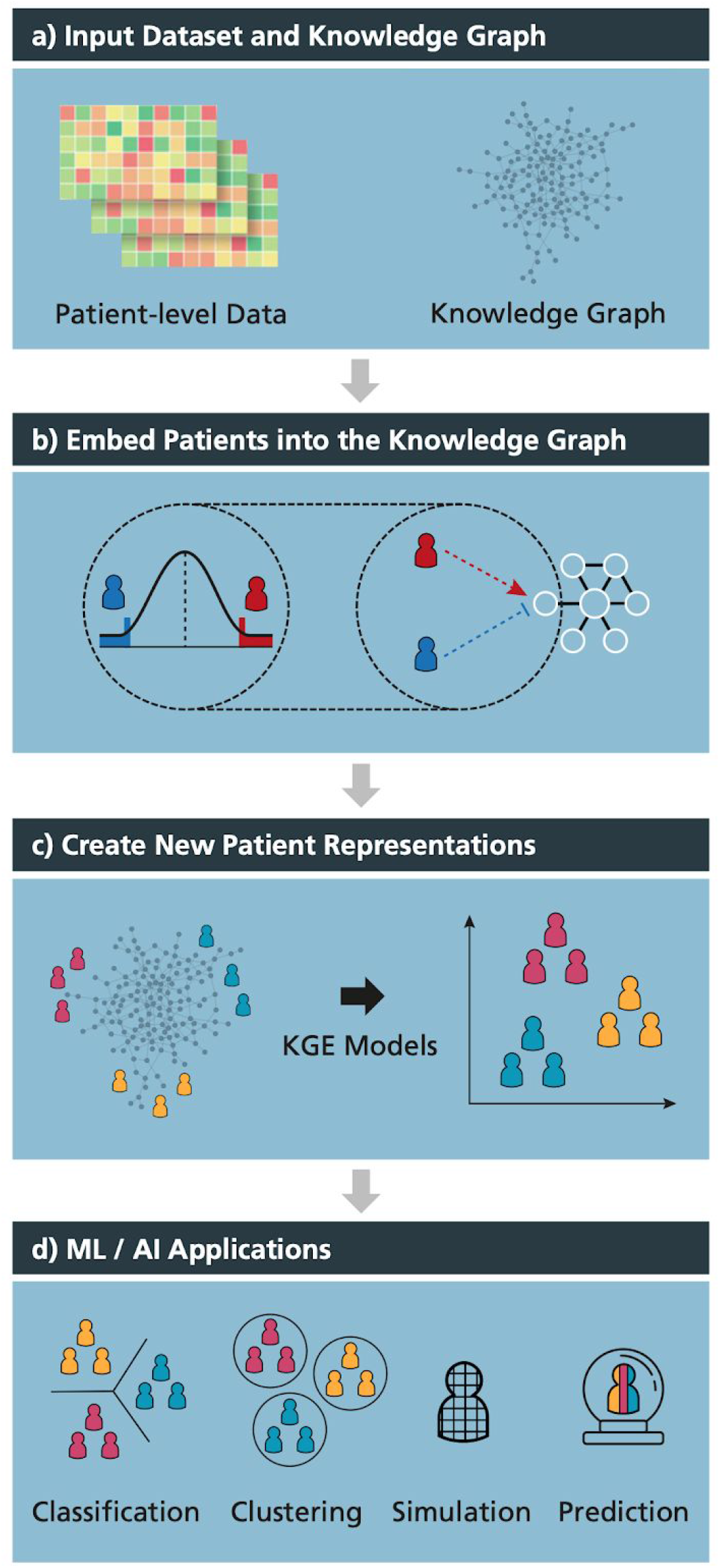
Schematic illustration of the framework. **a)** CLEP requires two inputs: i) a patient-level dataset such as multi-*omics*, and ii) a KG comprising relations between features measured in the previously-mentioned dataset. **b)** Using one of the proposed methods, CLEP incorporates patients into the KG by connecting them to their most distinctive features in the dataset. **c)** KGEMs are then used to generate new patient representations based on both data- and knowledge-driven features. **d)** These patient representations can subsequently be used for several downstream tasks, such as patient classification and stratification. A high quality version of this figure is available at https://doi.org/10.6084/m9.figshare.12834605.

#### 2.1.1. Input Data

Our framework requires two inputs: a patient-level dataset and a KG **(Figure 1a)**. It can be applied to any dataset and KG so long as the dataset features can be mapped to nodes in the KG. In other words, if we intend to use CLEP on a transcriptomics dataset such as RNA-Seq, the KG must contain relationships between genes/proteins that are mappable to the transcripts that have been measured. We would like to note that although the framework does not require that all features in the dataset are also present in the KG, it is recommended to maximize this overlap.

#### 2.1.2. Incorporating Patients into the Knowledge Graph

The first step of the methodology consists of incorporating patients as nodes in the KG in order to subsequently generate novel individualized feature representations for each patient (i.e., embeddings) through the use of KGEMs **(Figure 1b)**. These models generate the embeddings by exploiting the topology of the KG. Generating comprehensive embeddings requires an approach that can position a patient node in the KG according to that patient’s most informative features (i.e., features that distinguish them from patients who possess clinically different features). This way, patient nodes with similar features would be close together in a network and would thus have similar embeddings, while patients with dissimilar features would be farther away **(see example in Figure 3b)**. On the other hand, connecting every patient to a large number of nodes in the KG instead of a smaller node set that represents a patient’s most relevant features, would result in poor patient representations due to large overlaps of irrelevant features. In the methods and the supplementary sections, we describe numerous possible approaches designed to incorporate patients as nodes in the KG by identifying patient-specific features.

#### 2.1.3. Generating New Patient Representations

Once patients have been incorporated into the KG, the next step involves generating new patient representations through KGEMs **(Figure 1c)**. We restricted ourselves to KGEMs since they consider both directionality and edge types, and our KG contains several directed edge types. Incorporating edge types during the learning process can help the model to differentiate between node types (e.g.., patients and biological entities) and node sub-types (e.g., diseased patients, and controls). CLEP has adopted PyKEEN [19] as the KGEM-software due to its wide range of functionalities (e.g., a large number of KGEMs, hyperparameter optimization functionalities).

#### 2.1.4. Applications of Patient Representations

Once novel patient representations have been generated with CLEP, they can be used for a vast number of applications including classification and clustering tasks **(Figure 1d)**. Because our approach leverages prior knowledge of known interactions, we hypothesize that the use of these representations can yield superior performances on these tasks with respect to the use of the original patient-level datasets from which they were generated. Furthermore, our methodology facilitates biologically meaningful interpretations of patient-level data by positioning patients into different KG neighborhoods which may potentially correspond to biological processes that are characteristic for specific patient subgroups.

### 2.2. Case Scenarios

#### 2.2.1. CLEP’s Representations Outperform Raw Data in Classifying Cognitively Impaired Patients and Healthy Controls

Here, we present the results of our methodology and demonstrate how CLEP can improve the performance of ML models on patient classification tasks within the context of Alzheimer’s disease (AD). We incorporated ADNI patients [20] into a protein-protein interaction KG (i.e., PPI-KG) in order to generate novel patient representations using various KGEMs **(see Methods)**. We then compared the performance of several ML models in distinguishing between cognitively impaired patients from controls depending on whether the input data was the original transcriptomics data or novel patient representations. We summarize the performance of each of the five ML models in **Figure 2 (a and b).**

**Figure 2.**
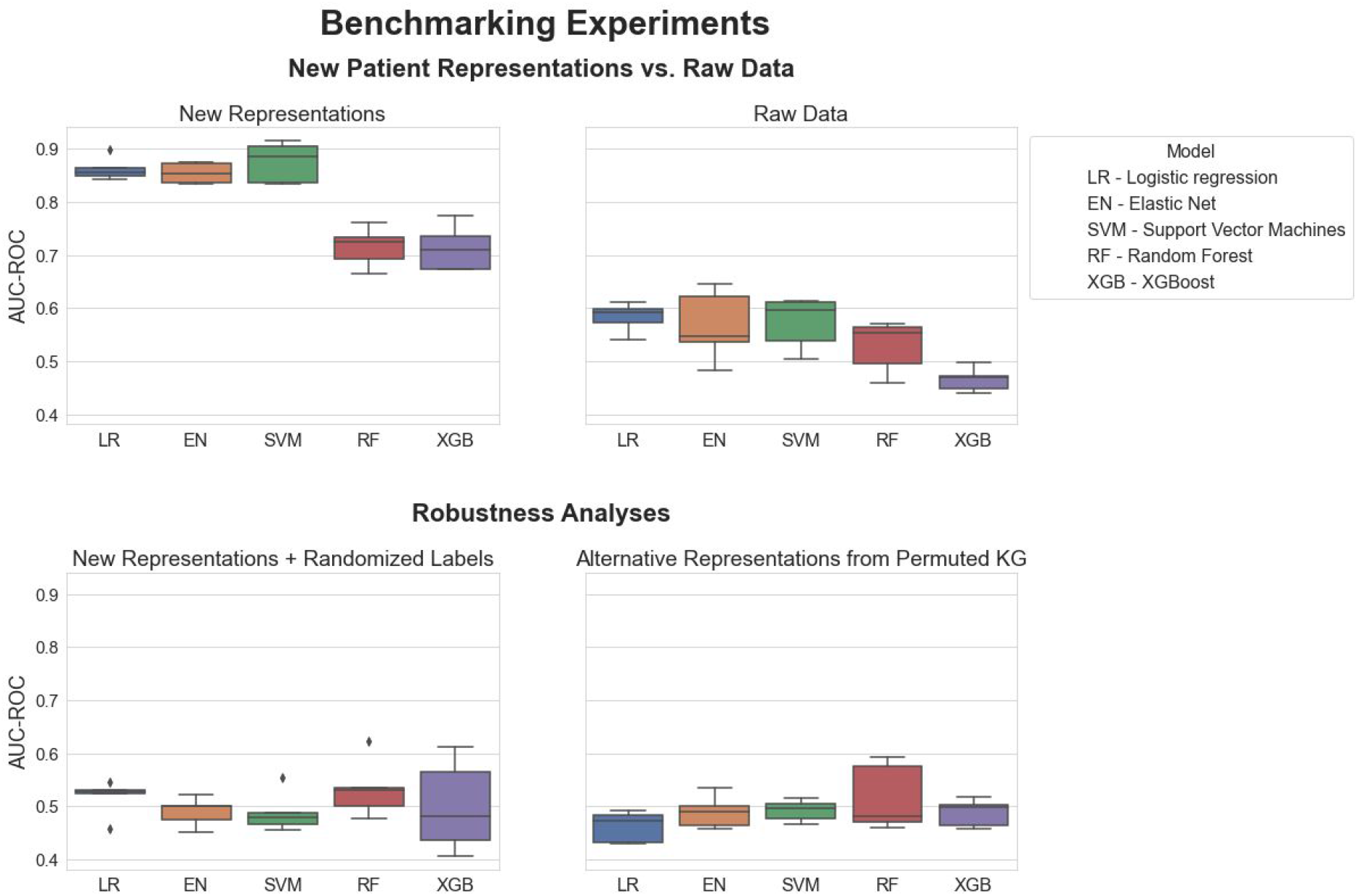
Benchmarking of five ML models trained to classify between cognitively impared patients and healthy controls. Each boxplot shows the distribution of the AUC-ROC values over 5 repeats of the 5-fold nested cross-validation procedure. Statistical modelling and ML methods are listed in **Table 1**. The new patient representations were generated by incorporating ADNI patients into the PPI-KG using a threshold of 2.5% on the eCDF of the control distribution for each mapped feature. The RotatE KGEM was trained on the KG using PyKEEN. The patient representations were used to train the five ML models with the original patient labels (a) and compared against the raw transcriptomics data (b). To investigate the robustness of our results, we randomized patient labels and trained the five ML models using the new representations (c). Furthermore, we trained the five ML models using the new representations generated from a permuted version of the original KG (d). Both robustness analyses yielded AUC-ROC values on all machine learning models equivalent to that of a random classifier (i.e., AUC-ROC values ~0.5).

**Table 1.** List of statistical modelling and machine learning methods available at CLEP for conducting classification tasks.

| Model | Reference | Implementation |
| --- | --- | --- |
| Logistic regression with $L_2$ regularization | - | scikit-learn [49] |
| Logistic regression with elastic net regularization | [45] | scikit-learn [49] |
| Support Vector Machines | [46] | scikit-learn [49] |
| Random Forest | [47] | scikit-learn [49] |
| XGBoost | [48] | XGBoost [48] |

Our results show that using original transcriptomics data as input for the binary classifier leads to relatively low prediction power (**Figure 2b)**, as opposed to the new representations generated by CLEP which substantially increase prediction performance (**Figure 2a)**. As an illustration, the SVM model, which was the highest performing, yielded area under the receiver operator characteristic curve (AUC-ROC) values ranging from 0.48 to 0.64 when trained on the transcriptomics data. In comparison, the same model, when trained on the new patient representations, yielded AUC-ROC values between 0.83 and 0.91. This difference was consistent across each of the five ML models, demonstrating how CLEP generates patient representations that significantly outperform the original data for this particular binary classification task. However, it is worth noting that the performance of the two tree-based methods (i.e., random forest and XGBoost) is significantly worse than the remaining models, yet they still outperformed their corresponding counterpart models trained on the raw data. Furthermore, to validate that the results are not an effect of the class imbalance between cognitively impaired patients and healthy controls [21], we reevaluated the ML models using the area under the precision-recall curve (AUC-PR) as a metric, which resulted in similar results **(Supplementary Figure 6)**. Finally, we would like to note that through the usage of the KGEMs, the dimension of the patient representations could significantly be compressed since the input dimension of the raw transcriptomics was larger than 40,000 features, while the representations generated by the KGEMs had a dimension of 256.

To investigate the robustness of our approach, we conducted two independent experiments. The first experiment consisted of training each of the five classifiers using new representations with randomized patient labels **(Figure 2c)** in order to confirm that the models did not fit to arbitrary artifacts in the data. In contrast, in the second experiment, patient representations were generated on a permuted version of the original KG **(see Methods for details)** to ensure that new representations reflect the information encoded in the KG by generating patient representations on a permuted version of the original KG **(Figure 2d)**. The results of these experiments yielded models with a performance equivalent to a random classifier (i.e., AUC-ROC values ~ 0.5); thus, confirming i) the robustness of our model evaluation strategy and ii) that the new representations are driven by information in the KG.

While we were able to show how CLEP successfully generated novel representations that yield superior prediction performance, the process of generating these representations is non-trivial. As an initial step, a threshold that determines which patient-measurable edges will be incorporated into the KG must first be selected as this parameter influences the KG generated **(see Subsection 4.1)**. We demonstrate the effect of the threshold on the previous binary classification task by training a single classifier using KGs derived from different thresholds **(Supplementary Figure 4)**. Unsurprisingly, while lower threshold settings (i.e., from 1% to 5%) resulted in increased predictive power, higher thresholds (i.e., 10% and 20%) penalized the performance of the model. This can be attributed to the fact that a low threshold exclusively creates connections between KG nodes and the patients falling at the extreme ends of a feature distribution, thus capturing patients that can best characterize a particular feature. On the other hand, higher thresholds generate a larger number of edges between patients and features, resulting both in a loss of specificity and the ability to distinguish between different patient groups.

The right choice of the KGEM is essential for generating patient representations that are useful for the classifiers. While the results of the case scenario are based on RotatE, we also generated patient representations based on further KGEMs **(Supplementary Figure 5)**. We can observe that patient representations generated by RotatE outperforms the remaining KGEMs by a large margin. It is known that specific relational patterns that can be modeled by RotatE cannot be modeled by TransE, ComplEx, and HolE. However, to obtain a clearer picture, relational patterns around patient nodes could be investigated. Finally, we also investigated the effect the size of a KG can have on results by evaluating the performance of ML models on smaller versions of the PPI-KG. As expected, smaller PPI-KGs resulted in lower performance for each of the ML models **(Supplementary Figure 7)**.

#### 2.2.2. Biological Interpretation and Patient Subgroup Identification through CLEP

Though the advent of machine learning methods have led to a new range of applications in the biomedical field, these methods inherently come with a tradeoff with regards to interpretability. So called black-box models lack transparency, providing no explanation as to what accounts for the predictions they generate. In the biomedical field in particular, this can often translate to a lack of success in clinical practice [1]; while discrete patterns that arise from ML techniques can be discerned, without an understanding of the fundamental cause and the characteristic features of a disease, clinicians may not have an adequate level of knowledge to make a confident diagnosis. In this section, we demonstrate how the hybrid KG generated by CLEP can drive the identification of patient-specific mechanisms and pathways made possible by the incorporation of patients into a KG comprising mechanistic knowledge **(Figure 3a)**.

**Figure 3.**
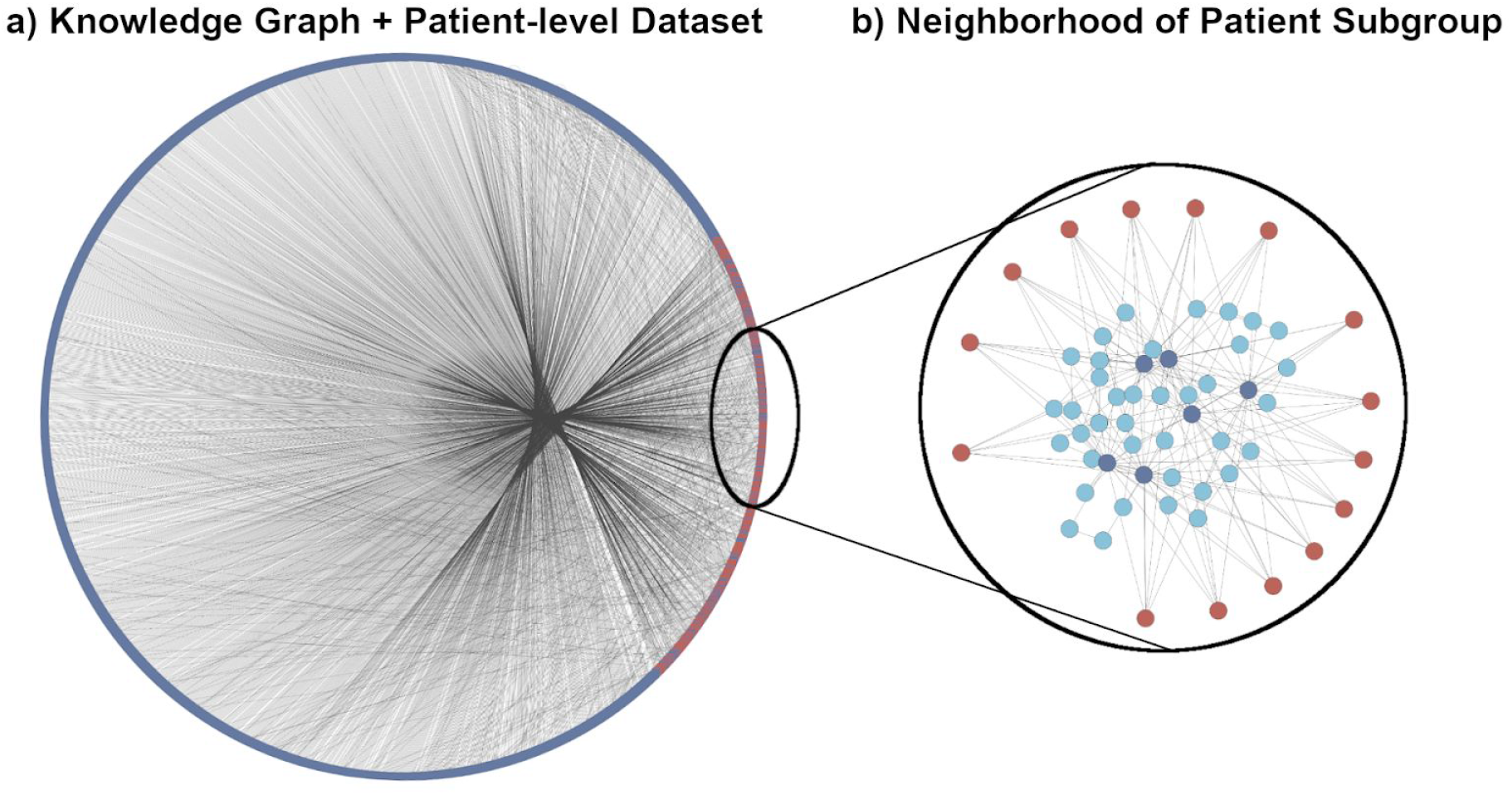
Incorporating patients into a KG fosters biological interpretation and identification of patient subgroups. **a)** PPI-KG subgraph after incorporating ADNI participants visualized using a circular layout. Red nodes represent ADNI patients and blue nodes represent proteins present in the PPI-KG. The majority of connections are between ADNI patients and their most characteristic proteins, which is represented by the large number of outgoing edges between the right part of the circle (where patients are located) and the rest of the KG. **b)** Local neighborhood around the subgroup of cognitively impaired patients investigated in the case scenario. Red nodes represent patients, dark blue the six proteins linked to the patient subgroup, and light blue other related proteins present in the neighborhood.

In the previous subsection, we demonstrated how novel representations generated on the ADNI dataset outperformed raw data in a diagnosis prediction task. However, these representations cannot be directly interpreted as they are low-dimensional vector representations embedded in a latent space. Nonetheless, because the original KG comprises biological knowledge, features and/or mechanisms that are connected to a given embedded patient or patient subgroup can be easily pinpointed by studying their local neighborhood in the KG. Thus, using the KG derived from the ADNI dataset, we identified sets of genes with connections to cognitively impaired patients (groups) but without any connections to control participants. Of these gene sets, we focused on a particular set of genes (HS2ST1, ESR2, IKBKG, UBE2D3, PCGF5, and NFIC), all of which were connected to each other, and were also identified in a subgroup of cognitively impaired patients (*n*=15). We then investigated the interplay of genes in the local neighborhood of the KG around this subset of patients in order to identify and deconvolute common pathways that could be responsible for their phenotypic make-up **(Figure 3b)**.

To identify the biological pathways these genes participate in, pathway enrichment analysis was run on this gene set **(Supplementary Table 2).** Of the nine enriched pathways we identified (*q*-value < 0.05), six were related to Toll-like receptor signaling (specifically, Toll-like receptor 4 signaling) due to the involvement of IKBKG and UBE2D3 in this pathway. Interestingly, this inflammatory pathway is often noted for its association to AD [22, 23], while UBE2D3 has been proposed as a potential biomarker for Alzheimer’s disease [24]. Furthermore, HS2ST1 is a member of the heparan sulfate biosynthetic enzyme family and responsible for the synthesis of heparan sulfate (HS), the latter of which is known to cause protein aggregation and lead to neurodegenerative disease [25]. Additionally, ESR2 polymorphisms have been linked to cognitive impairment and an increased risk for AD, predominantly in women [26, 27]. It is worth noting that by investigating patient subgroups characterized by particular features, we can also assess whether they share common alleles. Finally, the remaining two genes, HFIC and PCGF5, are both present in the enriched “Gene expression (transcription)” pathway, and are associated with the regulation of transcriptional factors.

#### 2.2.3. CLEP’s Representations Outperform Raw Data in Classifying Psychiatric Conditions and Healthy Controls

We also reproduced our methodology on an additional dataset containing transcriptomics experiments on psychiatric conditions and healthy controls **(see Methods)**. The results for this dataset resembled the ones observed in the previous case scenario **(Figure 4)**. The patient representations generated by CLEP outperformed the original data in the binary classification task by a large margin. By investigating the individual ML models, we observed that the difference in performance between the two tree-based methods (i.e., random forest and XGBoost) and the rest was smaller for this dataset. Similarly to the previous case scenario, SVM yielded the highest performance. However, the variability in performance for the elastic net, random forest, and XGBoost was significantly larger compared to the ADNI dataset.

**Figure 4.**
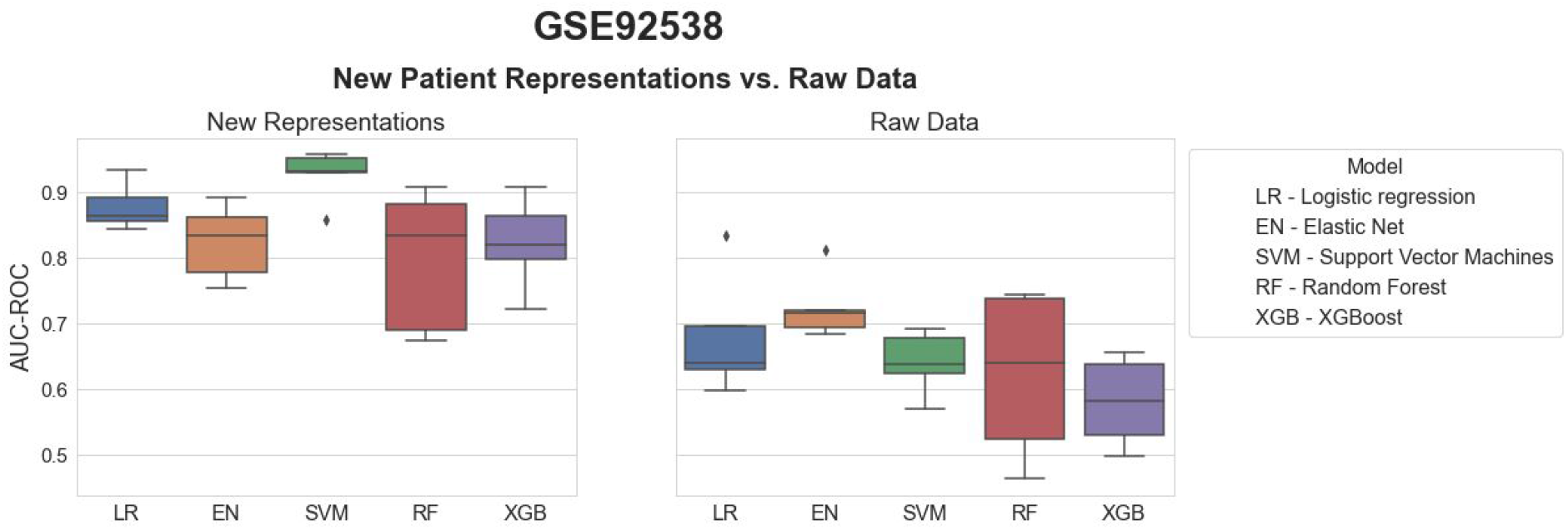
Benchmarking of five ML models trained to classify between psychiatric patients and healthy controls. Each boxplot shows the distribution of the AUC-ROC values over 5 repeats of the 5-fold nested cross-validation procedure. Statistical modelling and ML methods are listed in **Table 1**. The new patient representations were generated by incorporating the patien data from GSE92538 into the PPI-KG using a threshold of 2.5% on the eCDF of the control distribution for each mapped feature. The RotatE KGEM was trained on the KG using PyKEEN. The patient representations were used to train the five ML models with the original patient labels (a) and compared against the raw transcriptomics data (b).

## 3. Discussion

In this work, we have presented CLEP, a novel hybrid data- and knowledge-driven framework which leverages patient-level data and KGs for generating personalized patient representations. In the case scenarios, we demonstrated the utility of our framework on two independent datasets by employing transcriptomics data and an integrative KG containing knowledge from several protein-protein interaction databases. When compared to the raw transcriptomics data, we have shown how these representations yield superior performance in a binary classification task using a broad panel of machine-learning models. Furthermore, we have illustrated how incorporating knowledge from the KG for the generation of patient representations not only improves performance in classification tasks, but also facilitates the interpretation of the biological mechanisms that uniquely characterize patients and/or patient subgroups. In summary, we have shown the utility of a hybrid approach for the generation of new patient representations that can then be used in a broad range of applications, ranging from predictive modelling to personalized medicine. We have also made CLEP available as an extensible and reproducible software package, enabling researchers to conduct these experiments on various kinds of datasets and networks.

Our framework shares several limitations inherent to many ML methods. The first is the computational cost and time associated with training and optimizing KGEMs, which could be improved with the help of libraries focused on using multiple GPUs and distributing computation across a cluster. The second is that our methodology does not compensate for a lack of training data, as is often the case with clinical datasets, nor for poor quality data. The third is that this approach relies on the ability to meaningfully integrate two or more datasets with a KG. If there are too few mappable features between the clinical data and the KG, then the resulting patient representations may only have limited quality and utility, and some patients may be excluded entirely. Lastly, despite our framework being generalizable to any dataset, there may be cases where CLEP fails to improve the performance as the original dataset possesses sufficient informative power for a particular application. In these cases, however, one could always employ the KG to interpret and identify biological mechanisms (e.g., pathways in the KG) associated with an individual or group of patients, as demonstrated in our case scenario.

While we generated two types of edges (i.e., −1 or +1) between ADNI patients and proteins in the PPI-KG, we also implicitly generated negative edges between each ADNI patient and all other proteins to which neither a −1 nor +1 edge was inferred. Unfortunately, we were unable to explicitly incorporate them in the training algorithm because PyKEEN, as well as most KGEM packages, generate negative edges through uniform negative sampling or Bernoulli negative sampling [28] with respect to the given positive KG. Thus, we are interested to make improvements to the upstream package itself to enable these kinds of explicit inclusions, as there are a growing number of negative edges available in various biological knowledge sources.

There are a number of applications in precision medicine for the representations generated by CLEP. For instance, in the AD area, patient representations of cognitively impared patients could be systematically used to stratify patients in order to identify shared mechanisms that explain their observed phenotype. Furthermore, our approach can be generalized such that different types of patient-level data can be integrated and mapped to heterogeneous biological KGs, thus enabling scientists to combine their datasets with context-specific knowledge. In the future, we plan to extend this work as well as to adapt it to well-known network representation learning methods such as node2vec [29] and LINE [30]. Finally, we ambition that our framework could serve as an integration platform for multimodal datasets including clinical, imaging, and *-omics* data which could lead to more comprehensive patient representations.

## 4. Methods

### 4.1. Incorporating Patients into the Knowledge Graph

The method used to incorporate patients into the KG leverages the empirical cumulative distribution function (eCDF) **(Equation 1)**. The method first calculates the eCDF based on the quantitative measurements for each feature in the dataset which can be mapped to the KG. Using all samples in the dataset or exclusively the healthy controls data, we generate a reference distribution with data points that represent the measurements of each feature. We then identify patients that fall at the extreme ends of this distribution based on a predetermined threshold (e.g., 5% of the eCDF). Finally, these patients are connected to the node in the KG that represents that particular feature with an edge (i.e., −1 or +1) depending on the extreme in which the patient falls. The sign of the edge indicates whether a particular patient has a higher or lower measurement for that given feature to differentiate the patient from other patients in the distribution.

In the **Supplementary Text**, we present alternative methodologies that can similarly be used to incorporate patients to the KG. However, we have focused here on one specific method as it can be applied to any type of patient-level data and it does not make any assumptions on the feature distribution as opposed to other methods which either require pathway information or assume that the features are normally distributed.

### 4.2. Software Implementation

CLEP is implemented as a Python package to facilitate its usage within the scientific community. It contains several workflows corresponding to each of the steps presented in the methods from generating new patient representations to conducting downstream applications presented in the case scenario. Each workflow is both accessible through a command line interface (CLI) as well as programmatically, allowing users to input their own patient-level datasets and custom KGs. In total, CLEP offers three different methods for incorporating patients into the KG, all KGEMs available through PyKEEN [19], and five ML classifiers. Furthermore, thanks to its flexible implementation, users can independently use each of its modules as well as incorporate classifiers tasks into the framework **(Supplementary Figure 2)**. Finally, the source code of the CLEP Python package is available at https://github.com/hybrid-kg/clep under the Apache 2.0 License, its latest documentation can be found at https://clep.readthedocs.io, and it is distributed via PyPI at https://pypi.org/project/clep.

### 4.3. Case Scenarios

#### 4.3.1. Patient-level Data

The first dataset, the Alzheimer’s Disease Neuroimaging Initiative (ADNI) [20], is one of the world’s largest dementia cohorts and, with more than 1300 citations, the most referential resource for data-driven dementia research. In this work, we used the blood plasma transcriptomic data collected in the study (we refer to [31] and http://adni.loni.usc.edu for details). The dataset is already preprocessed and contains a total of 260 cognitively healthy control participants, 215 patients with early mild cognitive impairment, 225 patients with late mild cognitive impairment, and 44 patients with Alzheimer’s disease. To conduct the binary classification task, the latter three (i.e., all cognitively impaired patients) were grouped together into a single class (*n*=494). Preprocessed gene expression data (Robust Multichip Average (RMA) normalized data) was directly used as a baseline for the benchmarking of CLEP. To incorporate the patients into the KG, the expression of genes whose transcripts appear multiple times were considered individually.

The second dataset is a transcriptomics dataset containing samples from three psychiatric disorders (i.e., major depressive disorder, schizophrenia, and bipolar disorder) as healthy controls [32]. In total, this dataset contains 172 samples (41 major depressive disorder, 22 schizophrenia, 26 bipolar disorder, and 83 control samples). Similar to the ADNI dataset, the preprocessed gene expression data (RMA normalized data) from the second dataset was directly used as a baseline for the validation of CLEP. Similar to the previous dataset, we grouped the three psychiatric indications together to conduct a binary classification task.

#### 4.3.2. Knowledge Graph

For the case scenario, we employ a KG referred to as PPI-KG, which comprises protein-protein interactions from six resources: PathMe [33] (which includes KEGG [34], Reactome [35], and WikiPathways [36]), BioGrid [37], IntAct [38], and Pathway Commons [39] **(Supplementary Text)**.

#### 4.3.3. Generating Representations of Patients

We used different thresholds (i.e., the 1%, 1.5%, 2.5%, 5% 10%, and 20% quantiles of the eCDF) to define the tail of the distribution (i.e., extreme) which is required to incorporate ADNI patients and the patients of the GSE92538 dataset to the PPI-KG using the eCDF-based method described in **Subsection 4.1**. By applying this method on each threshold, we generated different KGs (i.e., one for each threshold) by connecting the patients that fall in the extremes of the reference distributions. Reference distributions were generated based on the expression values of normal patients for each gene that was present in both the transcriptomics data and the PPI-KG **(Supplementary Figure 3)**.

We would like to note that the resulting KG (i.e., PPI-KG after incorporating the ADNI patients) must be split into three triple sets (i.e., training, validation, and testing set) in order to train a KGEM. Thus, we developed an algorithm that splits a given KG in a way that relations and nodes are balanced across the three different splits. The pseudocode for this algorithm can be found in the **Supplementary Figure 1** and is implemented in the CLEP Python package.

#### 4.3.4. Selected Knowledge Graph Embedding Models

We selected RotatE [40], TransE [41], ComplEx [42], and HolE [43] to learn the patient embeddings, because they have been shown to be effective in a large-scale benchmarking study for KGEMs [44]. To train the final models, we used the best hyperparameters obtained by the hyperparameter optimization **(Supplementary Table 3).**

#### 4.3.5. Classifying between Cognitively Impaired and Healthy Controls

The new representations generated from KGEMs were used to classify between normal and cognitively impaired patients (i.e., AD and MCI) using five different statistical modelling and ML methods **(Table 1)**.

Prediction performance was evaluated via 5 times repeated 5-fold cross-validation in which the hyperparameters of the model were tuned within the cross-validation loop via a grid search **(Figure 5)**. The cross-validation prediction performance for the binary classification task was evaluated using the area under the receiver operating characteristic curve (AUC-ROC) or the area under the precision-recall curve (AUC-PR) as a metric.

**Figure 5.**
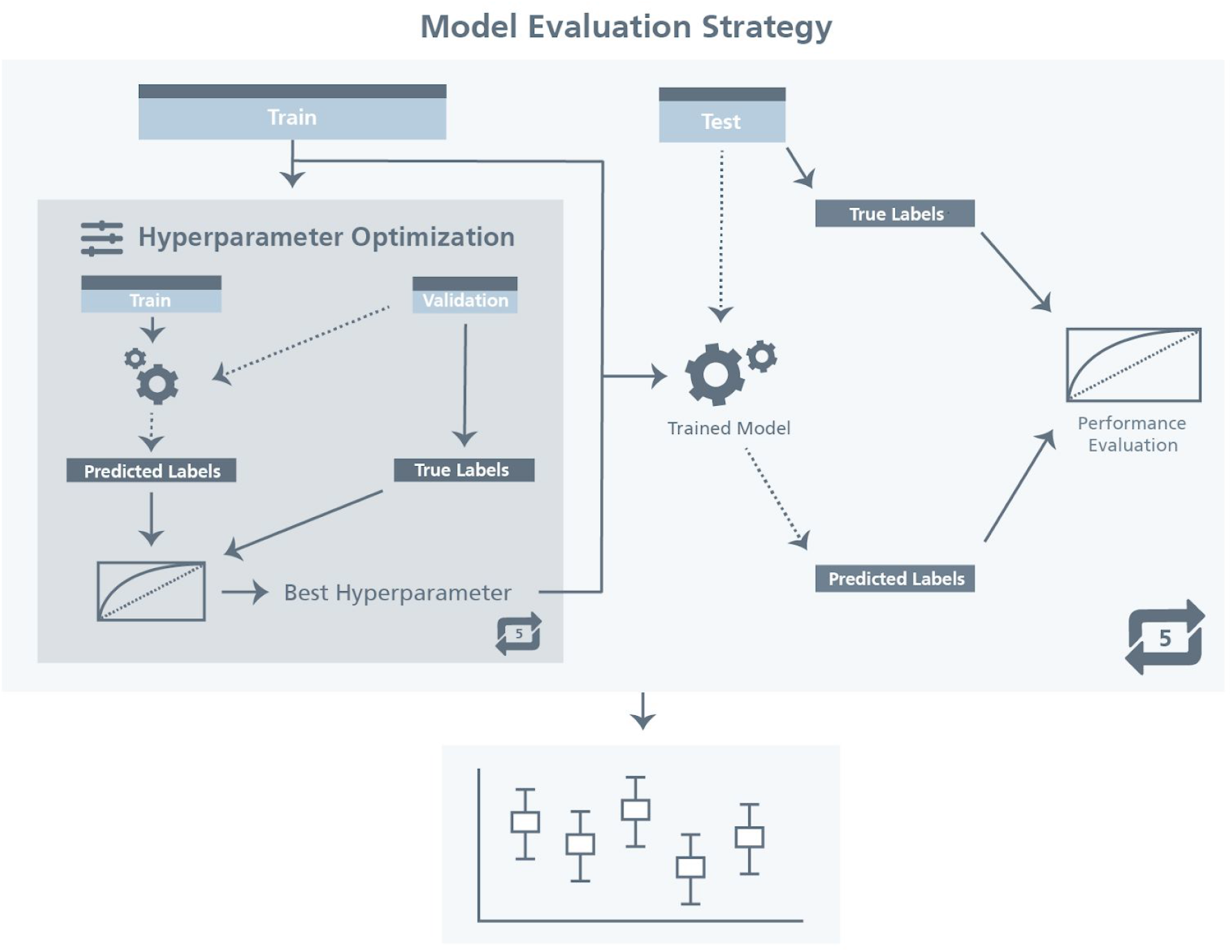
Schematic representation of the model evaluation strategy. The first step is to split the data into training (80%) and hold-out test set (20%). Next, we performed 5-fold stratified cross-validation by further repeatedly splitting the training data into 80% training and 20% validation in order to identify the best hyperparameter settings. Once these 5 cross validation rounds were performed, the best hyperparameters were used to train the model on the full training data and the trained model was evaluated on the held-out test set. This entire strategy was repeated 5 times to generate 5 AUC-ROC scores that were used to determine the performance of each model. We would like to note that the first split (training-test) ensured maintaining the overall class distribution and classes have been shuffled beforehand to avoid having the same data used in training/test. Furthermore, we chose AUC-ROC scores as evaluation metrics for both best hyperparameter selection and model evaluation. A high quality version of this figure is available at https://doi.org/10.6084/m9.figshare.12834608.

We then conducted three experiments to evaluate the robustness of our results. First, we trained the ML models using the new representations generated from RotatE with the 5% threshold while permuting patient labels (i.e., y-scrambling). Second, we trained the ML models on patient representations again generated using RotatE with a 5% threshold but using instead a permuted version of the KG generated using the XSwap algorithm [50] in order to investigate if a KG’s topology makes meaningful improvements to performance. By using this algorithm, we ensured that the permuted versions preserved the original structure of the original network (i.e., each node has the same number of in- and out-edges) but edges were randomly generated. Third, we trained the ML models using a subset of the PPI-KG corresponding to a single database (i.e., either WikiPathways or KEGG) in order to investigate the importance of KG size and completeness.

#### 4.3.6. Classifying between Psychiatric Disorders and Healthy Controls

Using the same setting used in the previous dataset **(Figure 5),** we trained the same five ML models to classify between normal samples and patients with a psychiatric disorder (i.e. bipolar disorder, major depressive disorder, and schizophrenia) using both the raw data and the new representations of the GSE92538 data generated by CLEP. Similar to the previous dataset, we used RotatE KGEM and a threshold value of 2.5% on either side of the control distribution for generating the new representations.

## Authors’ Contributions

DDF and CTH conceived and designed the study. VSB implemented CLEP and ran the experiments with supervision and support from DDF. MA guided and supported the implementation of the knowledge graph embedding generation and classification tasks. CB assisted with data and method development. SM processed the knowledge graph. MHA and JL acquired the funding. All the authors contributed to the writing of the manuscript.

All authors have read and approved the final manuscript.

## Acknowledgements

We would like to thank André Gemünd for his technical assistance running the experiments.

Data used in the preparation of this article were obtained from the Alzheimer’s Disease Neuroimaging Initiative (ADNI) database (adni.loni.usc.edu). As such, the investigators within the ADNI contributed to the design and implementation of ADNI and/or provided data but did not participate in analysis or writing of this report. A complete listing of ADNI investigators can be found at http://adni.loni.usc.edu/wp-content/uploads/how_to_apply/ADNI_Acknowledgement_List.pdf. Data collection and sharing for this project was funded by the Alzheimer’s Disease Neuroimaging Initiative (ADNI) (National Institutes of Health Grant U01 AG024904) and DOD ADNI (Department of Defense award number W81XWH-12-2-0012). ADNI is funded by the National Institute on Aging, the National Institute of Biomedical Imaging and Bioengineering, and through generous contributions from the following: AbbVie, Alzheimer’s Association; Alzheimer’s Drug Discovery Foundation; Araclon Biotech; BioClinica, Inc.; Biogen; Bristol-Myers Squibb Company; CereSpir, Inc.; Cogstate; Eisai Inc.; Elan Pharmaceuticals, Inc.; Eli Lilly and Company; EuroImmun; F. Hoffmann-La Roche Ltd and its affiliated company Genentech, Inc.; Fujirebio; GE Healthcare; IXICO Ltd.; Janssen Alzheimer Immunotherapy Research & Development, LLC.; Johnson & Johnson Pharmaceutical Research & Development LLC.; Lumosity; Lundbeck; Merck & Co., Inc.; Meso Scale Diagnostics, LLC.; NeuroRx Research; Neurotrack Technologies; Novartis Pharmaceuticals Corporation; Pfizer Inc.; Piramal Imaging; Servier; Takeda Pharmaceutical Company; and Transition Therapeutics. The Canadian Institutes of Health Research is providing funds to support ADNI clinical sites in Canada. Private sector contributions are facilitated by the Foundation for the National Institutes of Health (www.fnih.org). The grantee organization is the Northern California Institute for Research and Education, and the study is coordinated by the Alzheimer’s Therapeutic Research Institute at the University of Southern California. ADNI data are disseminated by the Laboratory for Neuro Imaging at the University of Southern California.

---

**Algorithm 1.**
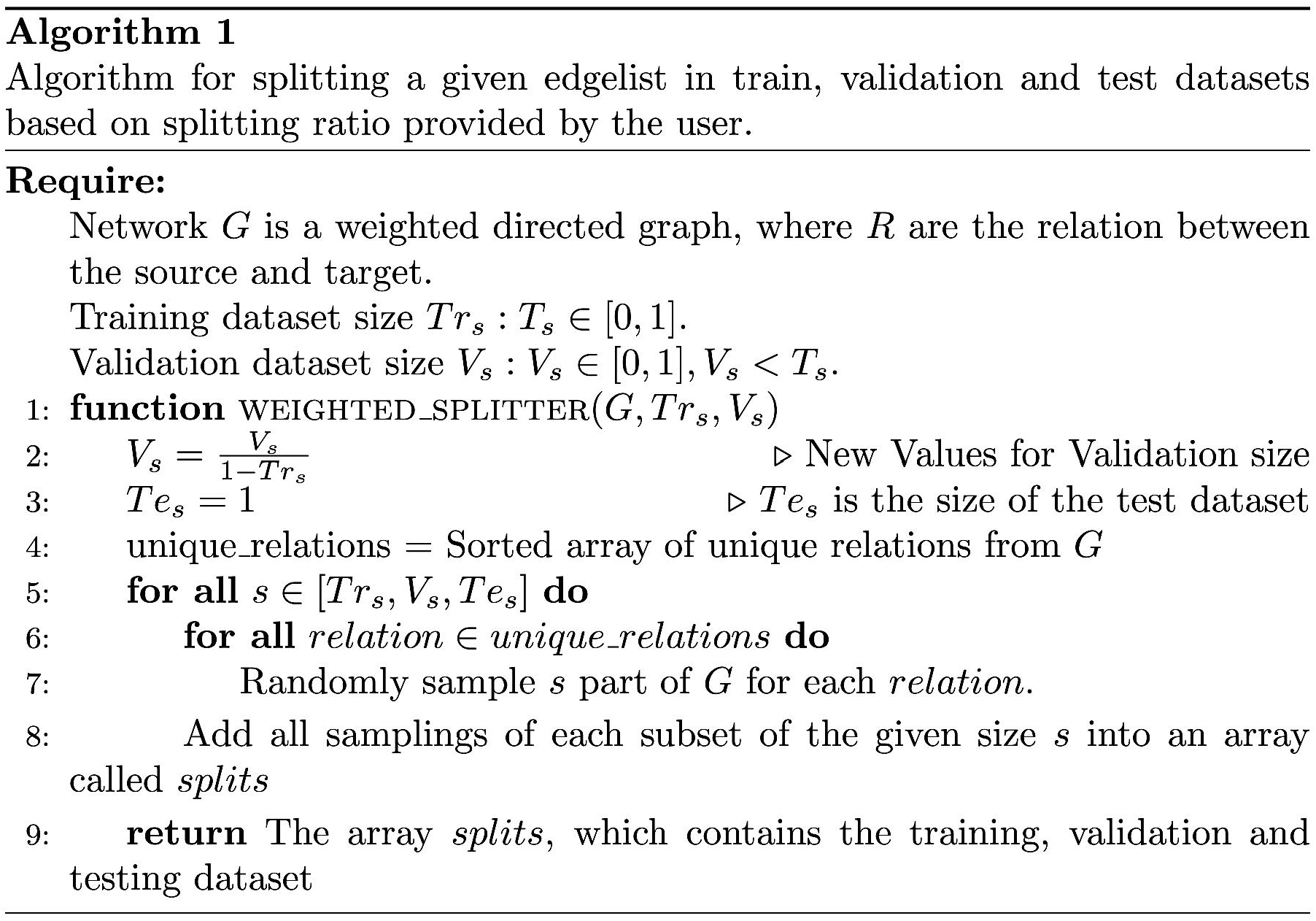
Algorithm for splitting a given edgelist in train, validation and test datasets based on splitting ratio provided by the user.

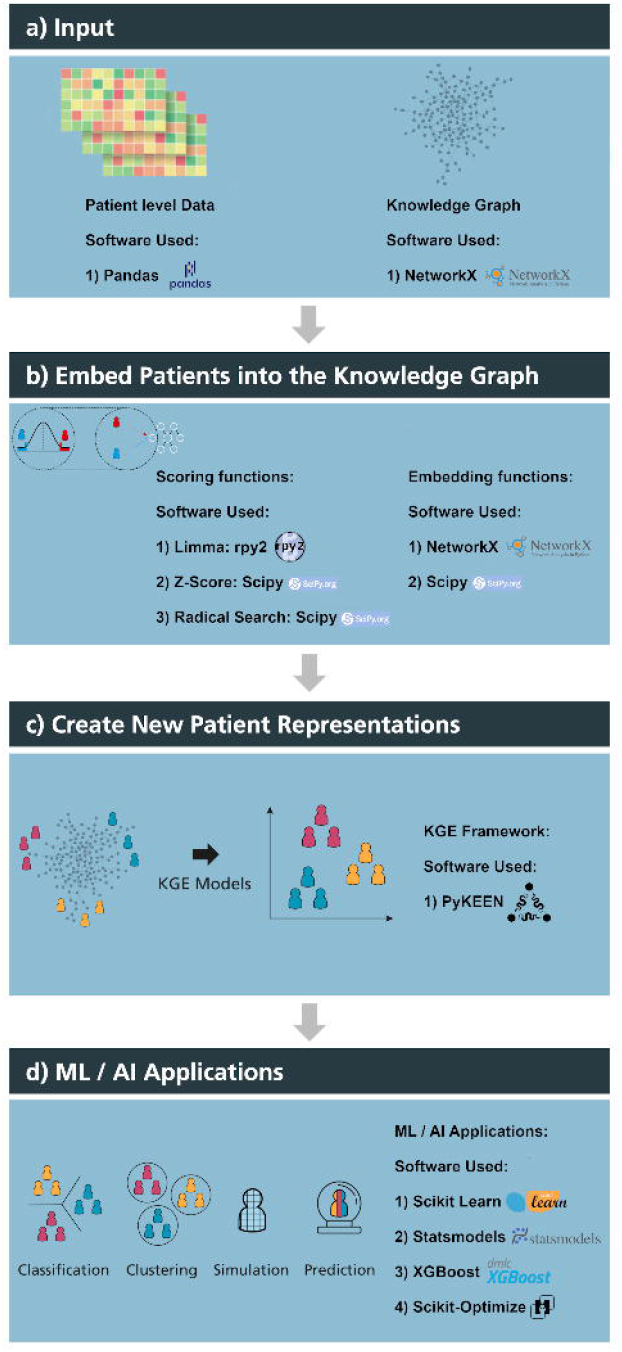

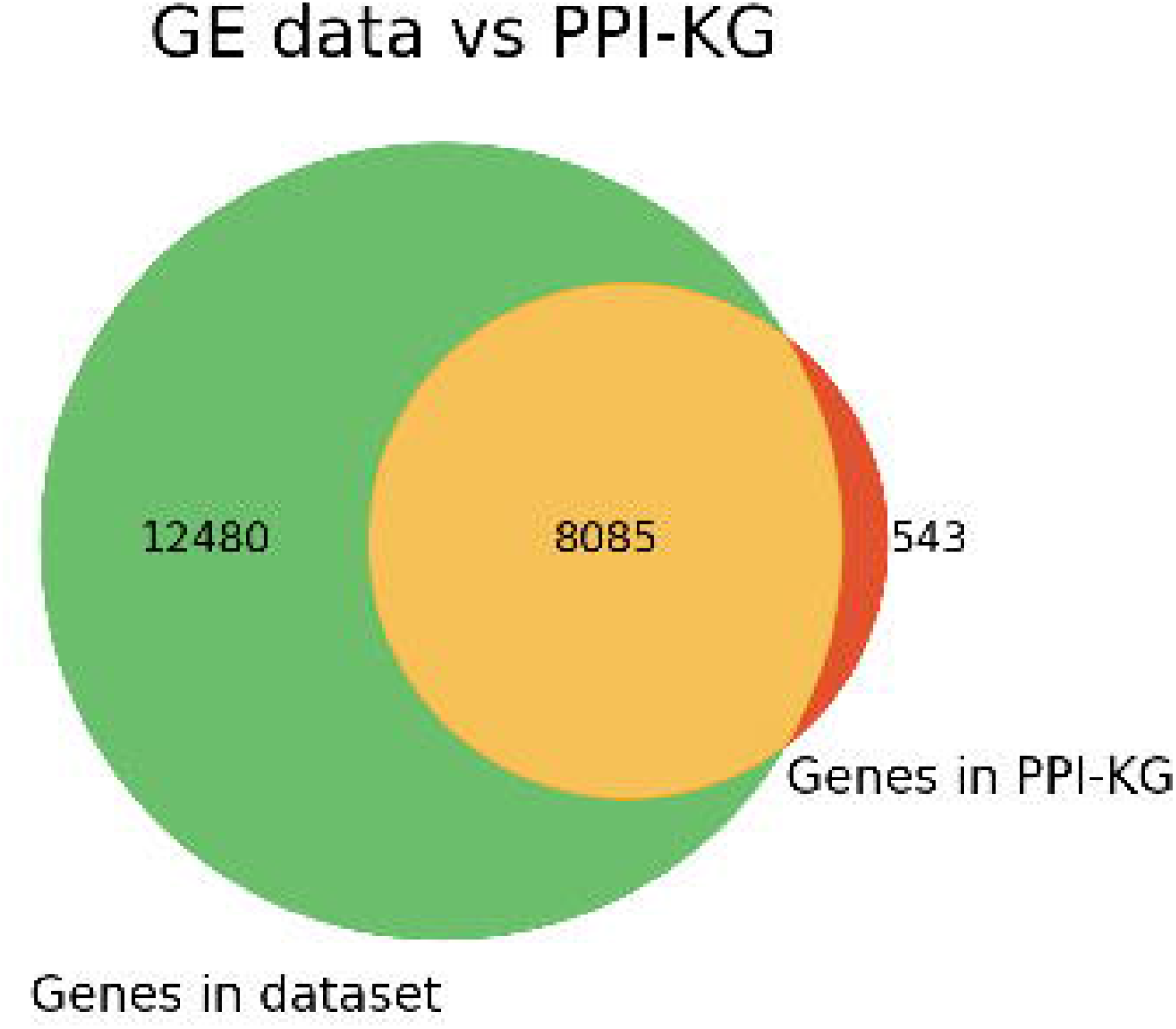

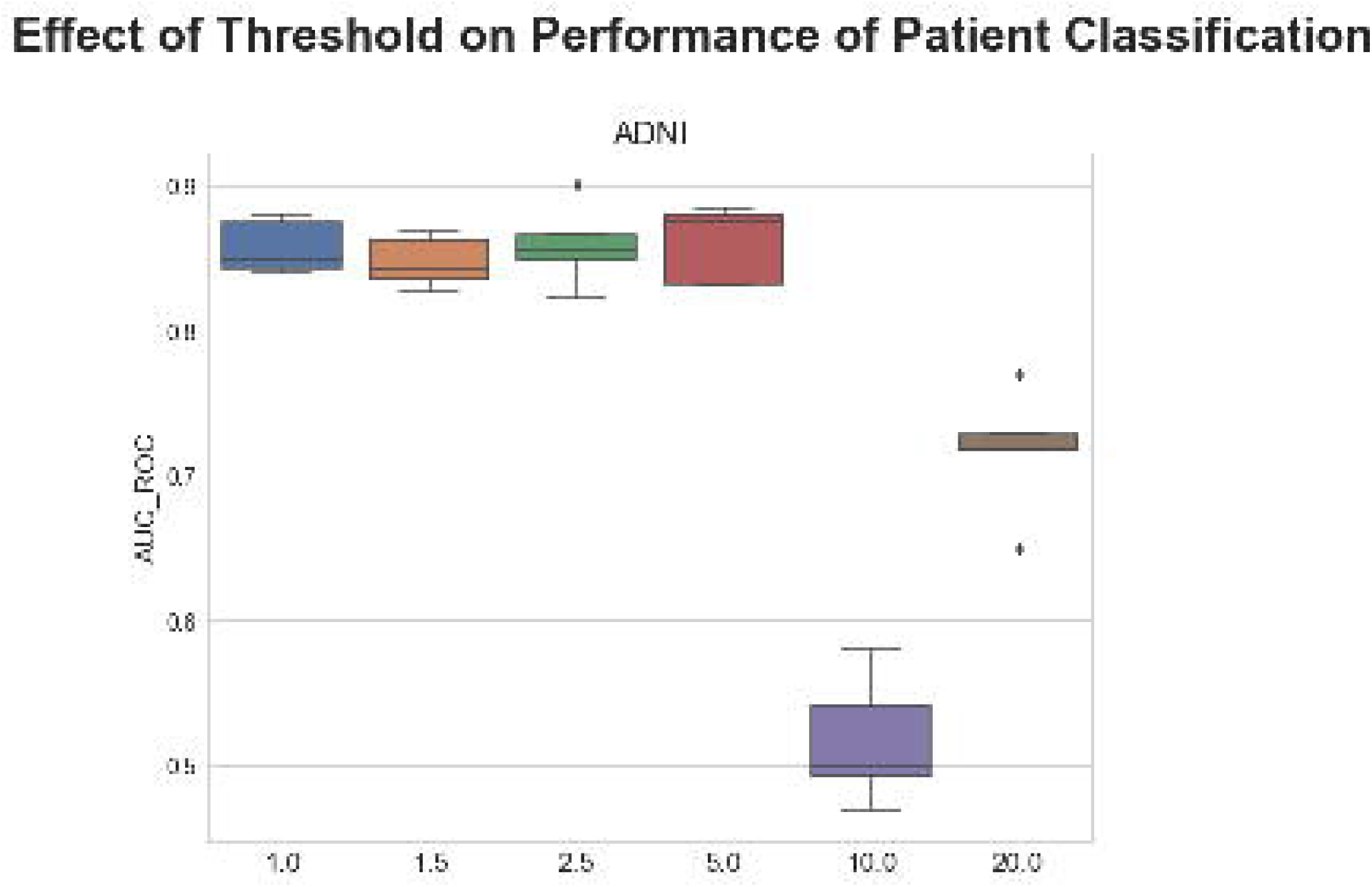

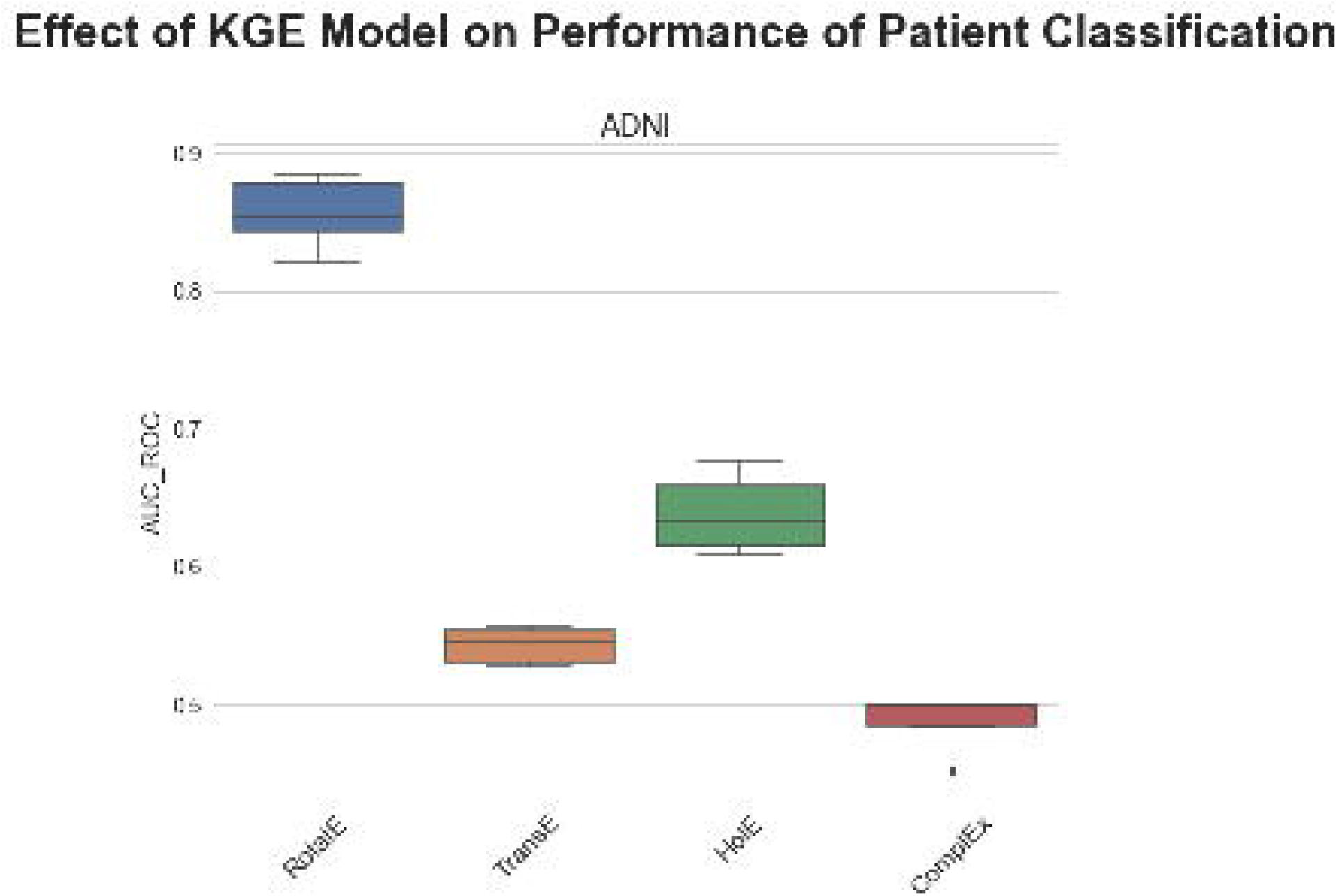

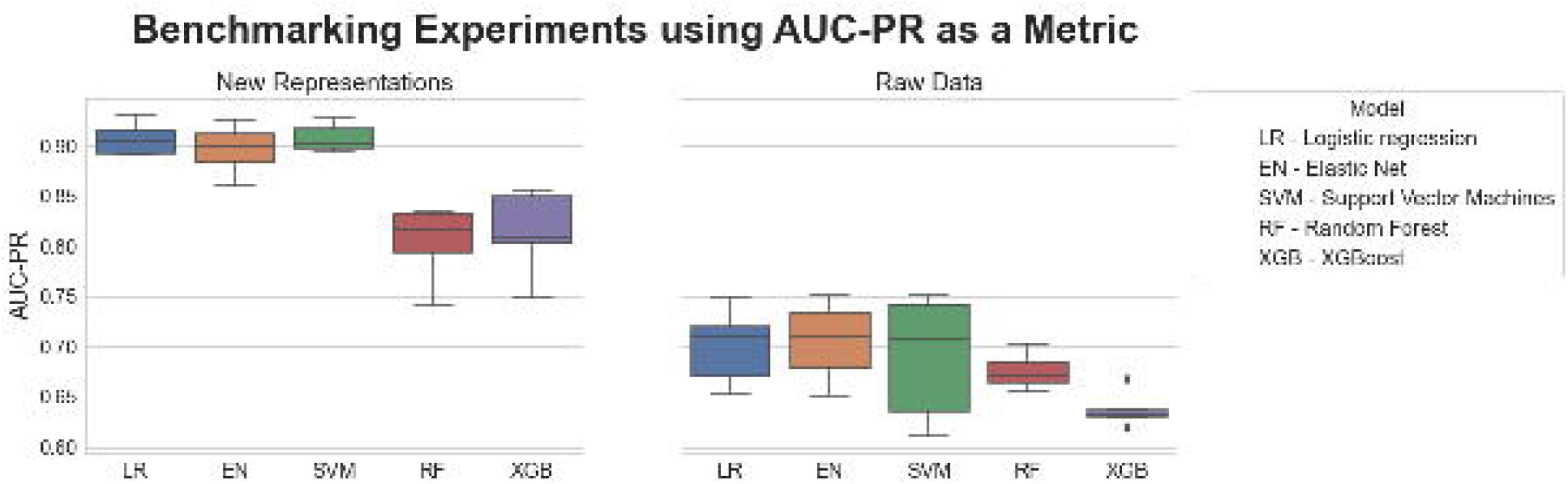

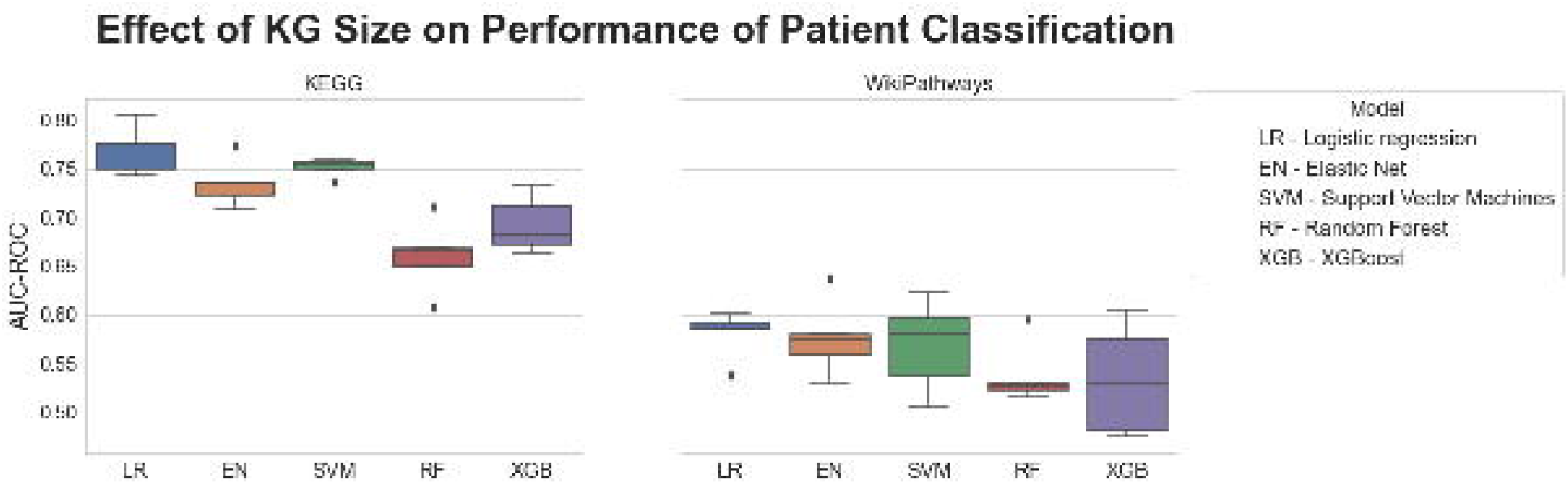

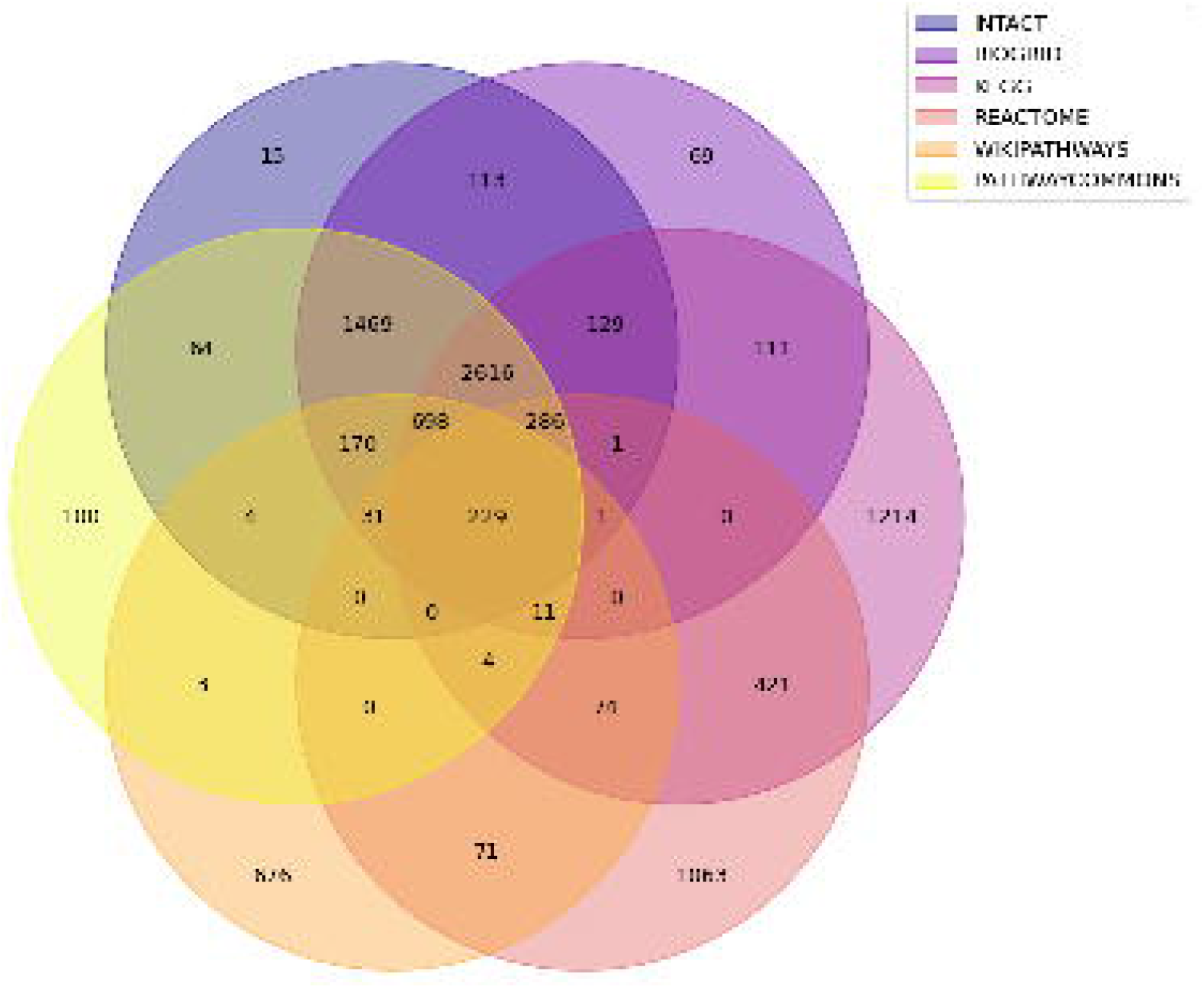

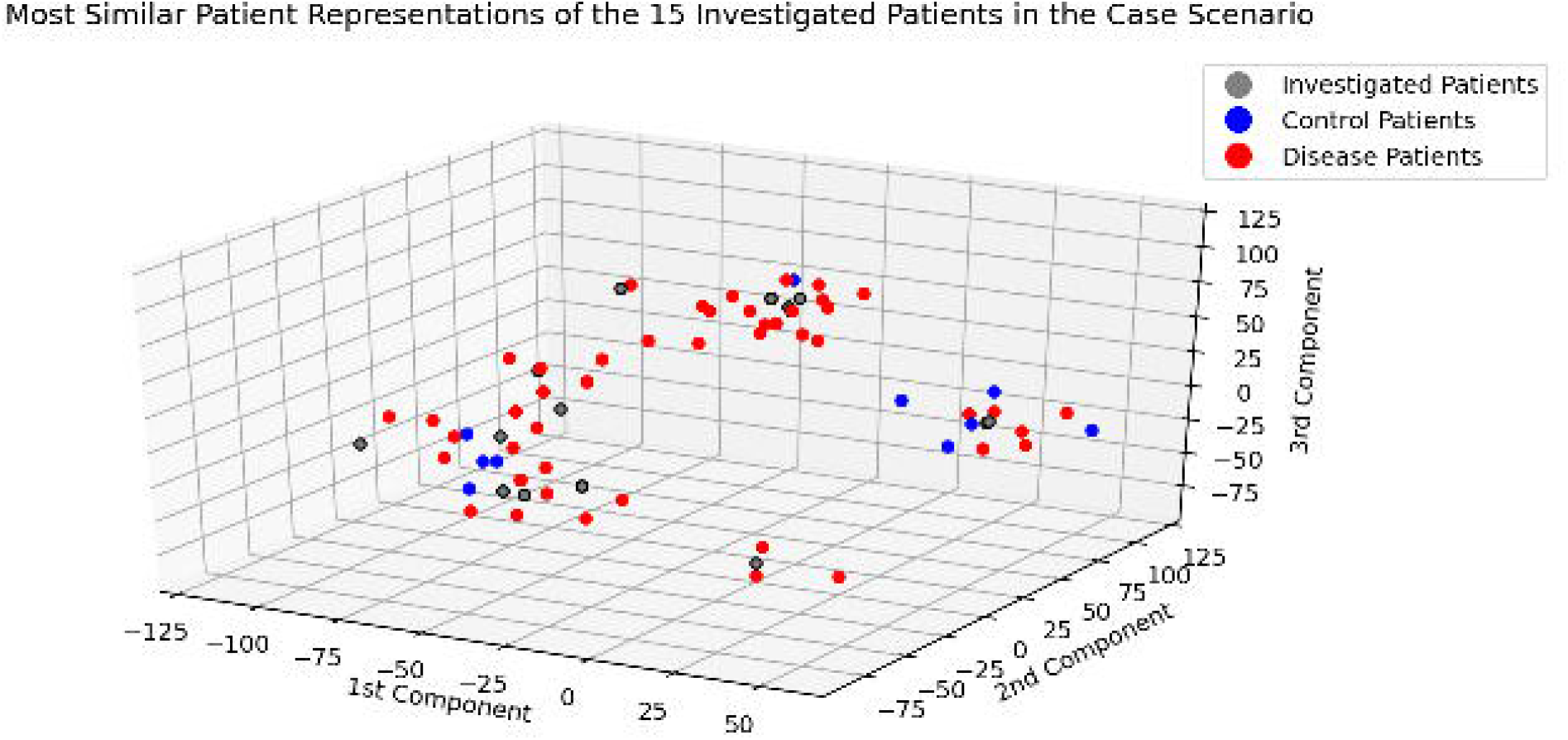

