## Supplementary File for "CLEP: A Hybrid Data- and Knowledge-Driven Framework for Generating Patient Representations"

### Additional File

#### Supplementary Text

##### 1. Additional Methodologies to Incorporate Patients into the KG

###### 1.1. Z-Score

The first of the alternative methods relies upon the measurement of the distance of a specific patient's feature with respect to a reference distribution of that feature. This reference distribution consists of all data points of that feature for a subset of the patients (e.g., control participants). Thus, this method requires to classify between at least two different patient groups. One important remark to consider is that this method assumes that the data is normally distributed. The method consists of the following steps:

1. For each feature in the dataset which can be mapped to the KG, the reference distribution is calculated using the data points from the reference participants.
2. Each patient is compared against this distribution using **Equation 1** and receives a patient score which measures the distance of that particular patient feature to the reference distribution.
3. Similar to the method presented in the paper, a given threshold determines the two extremes of the population and all patients that fall under the two extremes are connected with the node in the KG that represents the feature with a +1 or a -1 edge.

$$Z_i = \frac{x_i - \mu_{ref}}{\sigma_{ref}}$$

**Equation 1.** Z-score equation. *i* denotes a particular patient and *ref* denotes the data points present used as a reference distribution.

###### 1.2. Pathway Similarity Knowledge Graph

Pathway information is widely used in combination with *-omics* data both for interpretation and dimensionality reduction purposes. Thus, we propose an additional technique for KG generation that leverages pathway information. This technique called pathway-pathway overlap consists of annotating patient pathway scores (e.g., ssGSEA) to a KG representing pathway similarity. The first step, similar to other techniques we present, is to generate a KG that represents pathway similarity based on a similarity function (e.g., **Equation 2**). This function calculates the degree of overlap based on the entities on the pathway (e.g., gene set overlap) for each pair of pathways and only pairs that have a higher overlap than a predetermined threshold are connected with an edge on the KG. Once the similarity pathway KG has been generated, patients are connected to these pathways using the individual pathway scores for each patient given a predefined threshold. Thus, if the pathway score of a patient is significantly over-expressed (e.g., $\log_2$ fold change > 2.0 and $q$-value < 0.05), an up-regulation edge is created between the patient and the pathway, and vice versa if the pathway is under-expressed. Finally, this KG is used to generate the embeddings.

$$S_{(X,Y)} = \frac{|X \cap Y|}{min(|X|,|Y|)}$$

**Equation 2.** The Szymkiewicz-Simpson coefficient calculates the similarity between two sets (X and Y) where $0 \leq S \leq 1$. The similarity is the size of the intersection of the two sets divided by the size of the smaller set.

#### 2. Pseudocode for Generating the KG Train-Validation-Test Splits

**Algorithm 1**

Algorithm for splitting a given edgelist in train, validation and test datasets based on splitting ratio provided by the user.

**Require:**

Network $G$ is a weighted directed graph, where $R$ are the relation between the source and target.

Training dataset size $Tr_s : T_s \in [0, 1]$.

Validation dataset size $V_s : V_s \in [0, 1], V_s < T_s$.

**function** WEIGHTED_SPLITTER($G, Tr_s, V_s$)

  $V_s = \frac{V_s}{1-Tr_s}$ ▷ New Values for Validation size

  $Te_s = 1$ ▷ $Te_s$ is the size of the test dataset

  unique_relations = Sorted array of unique relations from $G$

  **for all** $s \in [Tr_s, V_s, Te_s]$ **do**

    **for all** $relation \in unique\_relations$ **do**

      Randomly sample $s$ part of $G$ for each $relation$.

    Add all samplings of each subset of the given size $s$ into an array called *splits*

  **return** The array *splits*, which contains the training, validation and testing dataset

**Supplementary Figure 1.** Pseudocode of the algorithm that generates distributed train-validation-test splits. The source code is implemented in Python and is available in CLEP at https://github.com/hybrid-kg/clep/blob/master/src/clep/embedding/kge.py.

#### 3. PPI-KG Generation

The PPI-KG was generated by concatenating each of the resources outlined in the paper using the command line interface of Bio2BEL (https://github.com/bio2bel) and PathMe (https://github.com/PathwayMerger/PathMe). Non-protein nodes were filtered since some of the resources contain other node types such as metabolites and miRNA. The relations present in the KG are encoded using Biological Expression Language (BEL) **(Supplementary Table 1)**. The PPI-KG is available at https://github.com/hybrid-kg/clep-resources/blob/master/Datasets/ADNI/kge_model/data/ppi-kg.edgelist.

| Relation Type | Frequency |
|---|---|
| Association | 152,433 |
| Increases | 22,750 |
| Decreases | 7,331 |
| Regulates | 21,023 |
| hasComponent | 6,098 |

**Supplementary Table 1. Frequency of relation types in the PPI-KG.**

#### 4. Hardware

KGEMs were trained on a GPU node with two Intel Xeon Scalable Gold 6140 processors per node with 18 cores/36 threads each (36 cores/72 threads per node), 2.3GHz base / 3.7 GHz Turbo Frequency, 768GB RAM (DDR4 ECC Reg) and eight NVIDIA Tesla V100 SXM2 32GB RAM GPUs (NVIDIA Volta Generation, 5120 CUDA cores, 640 Tensor cores, 32GB HBM2 memory, 300GB/s NVIDIA NVLink interconnect). The network was 100GBit/s Intel OmniPath, storage was 4x 3.2TB Samsung NVMe SSD PM1725a for local intermediate data and BeeGFS parallel file system for Home directories.

The training and evaluation of ML models were performed on a symmetric multiprocessing (SMP) node with four Intel Xeon Platinum 8160 processors per node with 24 cores/48 threads each (96 cores/192 threads per node in total) and 2.1GHz base / 3.7 GHz Turbo Frequency with 1536GB/1.5TB RAM (DDR4 ECC Reg). The network was 100GBit/s Intel OmniPath, storage was 2x Intel P4600 1.6TB U.2 PCIe NVMe for local intermediate data and BeeGFS parallel file system for Home directories.

### Supplementary Figures

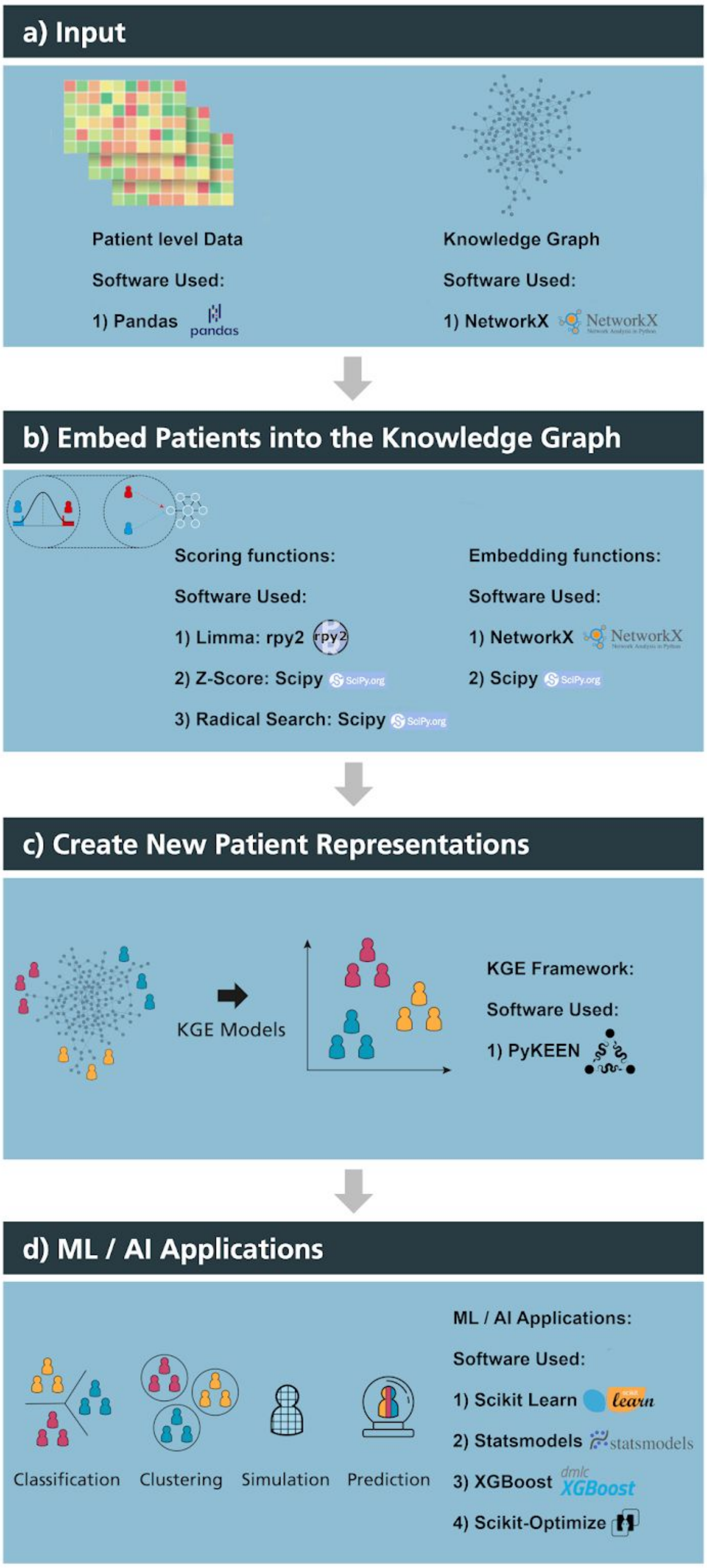


**Supplementary Figure 2. Libraries employed by CLEP.** CLEP uses Pandas to store the patient-level data and NetworkX to store the KG as a network. The scoring method *limma* utilises the rpy2 library while the others use SciPy. For training KGEMs, CLEP leverages the PyKEEN library. Finally, ML/AI applications use various statistical Python libraries such as Scikit-learn, statsmodels, Scikit-Optimize, and XGBoost.

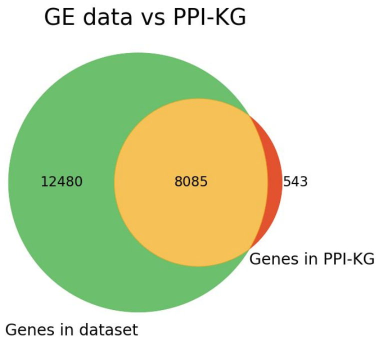


**Supplementary Figure 3. Venn diagram displaying the overlap between features in the ADNI transcriptomic dataset and proteins in the PPI-KG.** 8,085 of the total 20,565 genes measured in the transcriptomics dataset can be mapped to their corresponding proteins in the PPI-KG.

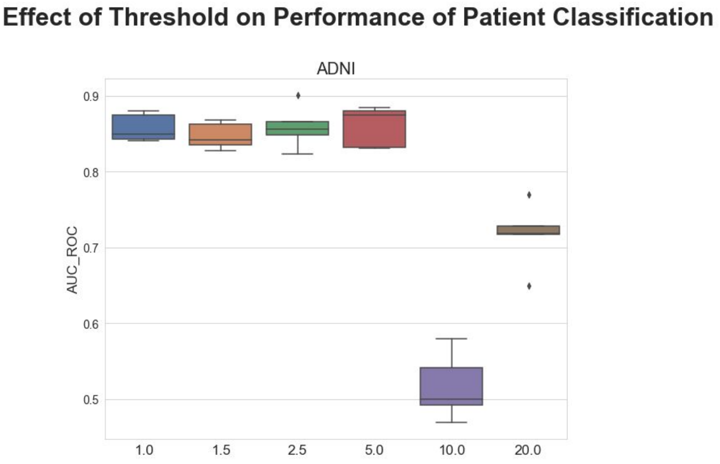


**Supplementary Figure 4. Effect of the chosen threshold on the performance of the elastic net regression model.** Each boxplot shows the distribution of the AUC-ROCs for the elastic net regression model over 5 repeats of the 5-fold nested cross-validation procedure across varying thresholds (y-axis). As expected, while low thresholds (i.e., lower than 5%) yield new patient representations with high prediction power, higher thresholds (i.e., 10% and 20%) yield lower performance as the number of connections for each patient increases. Thus, patients are not only connected to their most characteristic features but to a greater number of features which ultimately hinders the generation of comprehensive representations by the KGEM. The number of connections between patients and genes for each threshold is shown in **Supplementary Table 4**.

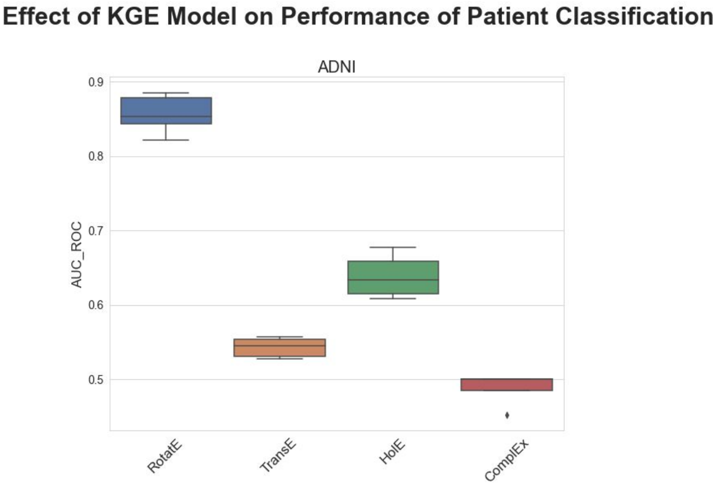

**Supplementary Figure 5. Comparison of different KGEMs on the performance of the elastic net regression model.** Each boxplot (i.e., elastic net regression model) shows the distribution of the AUC-ROCs over 5 repeats of the 5-fold nested cross-validation procedure for varying KGEMs (y-axis).

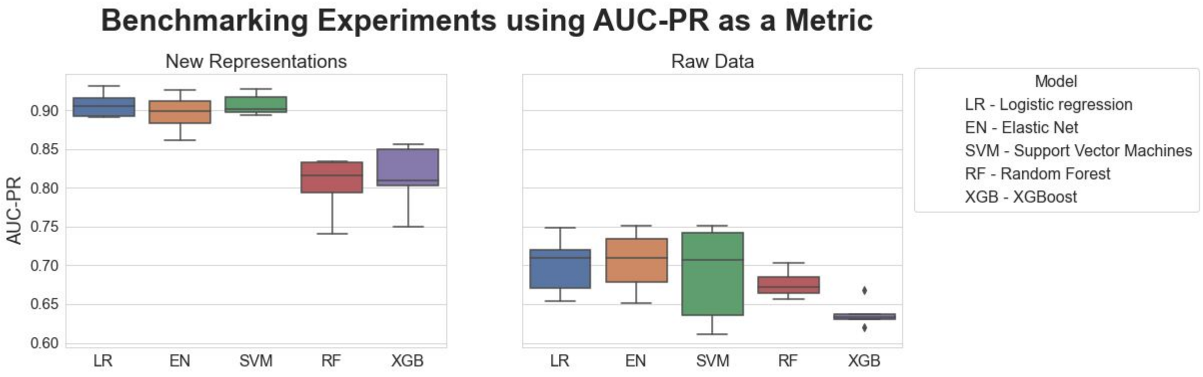


**Supplementary Figure 6. Benchmarking of five ML models trained to classify between cognitively impaired patients and healthy controls.** Each boxplot shows the distribution of the area under the precision-recall curve (AUC-PR) values over 5 repeats of the 5-fold nested cross-validation procedure. The new patient representations were generated by incorporating ADNI patients into the PPI-KG using a threshold of 2.5% on the eCDF of the control distribution for each mapped feature. The resulting KG was trained using the RotatE KGEM on PyKEEN. The new patient representations generated with CLEP were used to train the five ML models (left) and compared against the raw transcriptomics data (right).

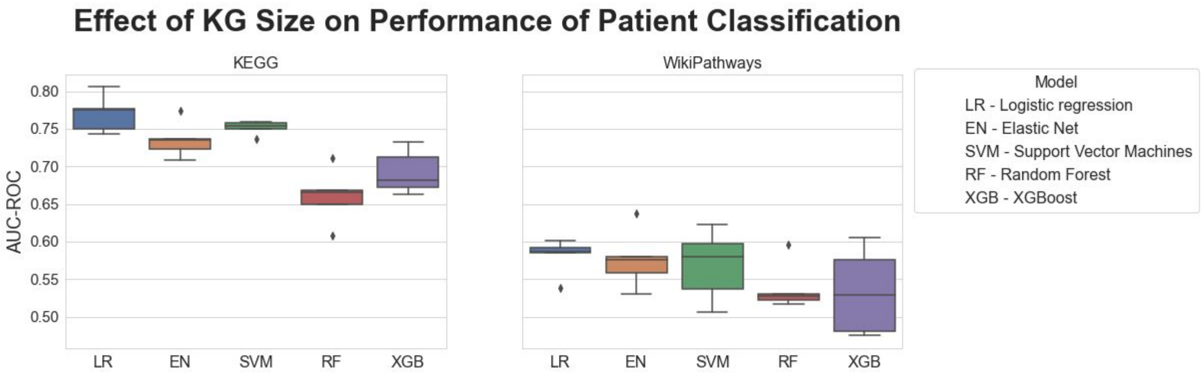


**Supplementary Figure 7. Benchmarking of five ML models trained to classify between cognitively impaired patients and healthy controls on the KEGG and WikiPathways subgraphs of the PPI-KG.** Each boxplot shows the distribution of the AUC-ROCs over 5 repeats of the 5-fold nested cross-validation procedure. The new patient representations were generated by incorporating ADNI patients into the KEGG/WikiPathways subgraphs of the PPI-KG using a threshold of 2.5% on the eCDF of the control distribution for each mapped feature. The resulting KG was trained using the RotatE KGEM on PyKEEN.

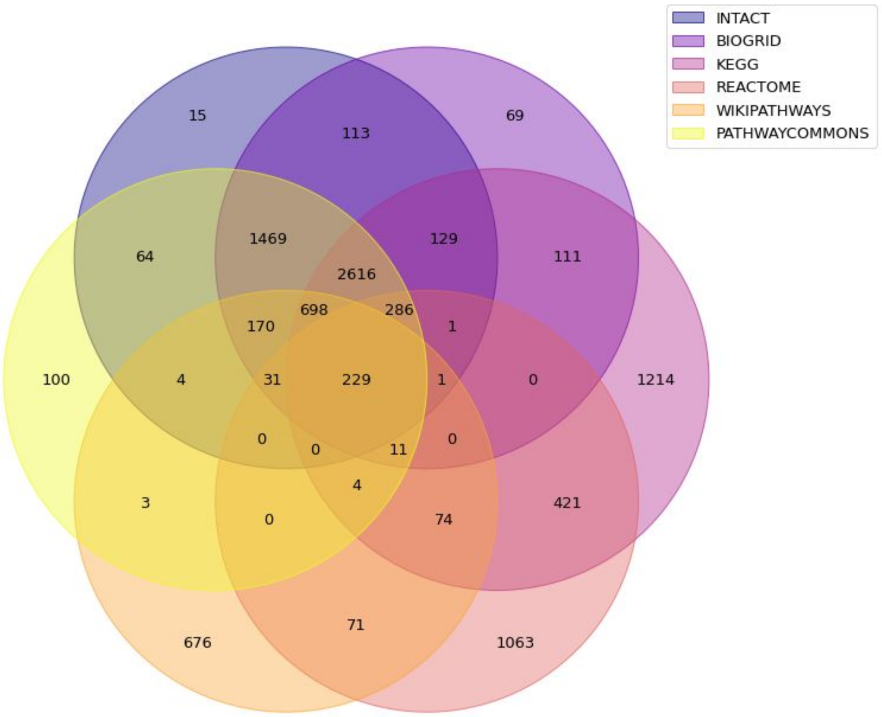


**Supplementary Figure 8. Venn diagram displaying the overlap between the different datasets in the PPI-KG.**

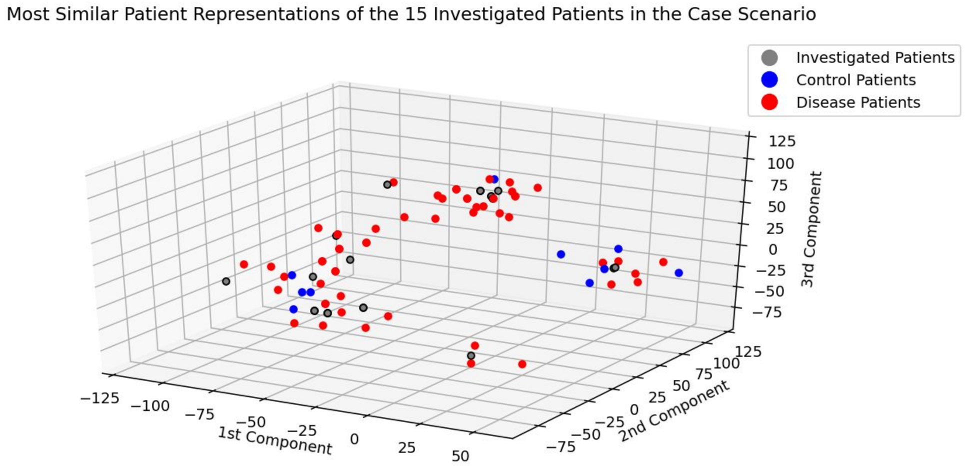


**Supplementary Figure 9. Visualization of the nearest neighbors to the 15 investigated patients in the case scenario using t-SNE.** The plot depicts the 15 investigated patients and their six nearest neighbors using the three first components of t-SNE. The 15 investigated patients are colored in grey, control patients in blue and cognitively impaired patients in red.

### Supplementary Tables

| **Database** | **Pathway** | **Identifier** | ***q*-value** |
|---|---|---|---|
| Reactome | Toll-Like Receptor 4 (TLR4) Cascade | R-HSA-166016 | 0.043411 |
| Reactome | DDX58/IFIH1-mediated induction of interferon-alpha/beta | R-HSA-168928 | 0.028475 |
| Reactome | MyD88-independent TLR4 cascade | R-HSA-166166 | 0.028475 |

| | | | |
|---|---|---|---|
| Reactome | TRIF(TICAM1)-mediated TLR4 signaling | R-HSA-937061 | 0.028475 |
| Reactome | Toll-Like Receptor 3 (TLR3) Cascade | R-HSA-168164 | 0.028475 |
| WikiPathways | DNA IR-damage and cellular response via ATR | WP4016 | 0.028475 |
| Reactome | Gene expression (Transcription) | R-HSA-74160 | 0.021049 |
| Reactome | IKK complex recruitment mediated by RIP1 | R-HSA-937041 | 0.006218 |
| Reactome | TICAM1, RIP1-mediated IKK complex recruitment | R-HSA-168927 | 0.006218 |

**Supplementary Table 2. Enriched pathways (*q*-value < 0.05) after running over-representation analysis on the gene set studied in the case scenario.** Enrichment analysis was conducted using pathways from the Reactome and WikiPathways databases downloaded on 16-08-2020.

| Hyperparameter | Best Obtained Value |
|---|---|
| Embedding Dimension | 128 |
| Training Approach | Stochastic Local Closed World Assumption |
| Number of negatives per positive | 22 |
| Loss Function | Negative Sampling Self-Adversarial Loss |
| Margin | 23.92 |
| Adversarial Temperature | 0.93 |
| Batch Size | 1024 |
| Number of Epochs | 200 |

**Supplementary Table 3. Best hyperparameters for training the RotatE model on PPI-KG + ADNI.**

| Threshold | Number of Patient-Feature Edges |
|---:|---:|
| 1.0 | 726,566 |
| 1.5 | 1,040,551 |
| 2.5 | 1,732,189 |
| 5.0 | 3,342,029 |
| 10.0 | 6,363,445 |
| 20,0 | 12,055,826 |

**Supplementary Table 4. Number of edges between patients and mapped features in the KG for each of the thresholds employed.**
